# Cell Trajectory Inference based on Schrödinger Problem and a Mechanistic Model of Stochastic Gene Expression

**DOI:** 10.1101/2025.11.21.689689

**Authors:** Clémence Fournié, Elias Ventre, Ulysse Herbach, Aymeric Baradat, Olivier Gandrillon, Fabien Crauste

## Abstract

Cellular differentiation is the biological process that leads a cell to opt for a particular cellular identity. Recently, single-cell RNA-sequencing has enabled the simultaneous measurement of gene expression levels at specific times for a large number of individual cells and a large number of genes. Repeating such measurements at different time points gives then access to the temporal variation, or transport, of a distribution on a gene expression space. The whole temporal trajectory of distributions thus characterizes the differentiation process at population level, but trajectories of individual cells are still out of reach since most measurement techniques are destructive.

The optimal transport theory that has been used so far to infer cellular differentiation trajectories from time-stamped single-cell RNA-seq data involves solving the so-called Schrödinger problem in its most common version. This implies assuming that cells move, in the gene expression space, by diffusion. Yet, real gene dynamics are much more complex.

In the present work, we assume that mRNA dynamics are characterized by brief and important production of RNA, with long periods of inactivity in between, and consider the so-called Bursty model of gene dynamics. We use this model to define a reference process for the Schrödinger problem. By comparing the solutions of the Schrödinger problems with a Diffusive and a Bursty reference process, under different conditions, we show that the Bursty model provides a better approximation of the underlying gene dynamics than the standard Diffusive process when inferring cell trajectories.

## 1 Introduction

While two cells in a multicellular organism may possess the same genome, they can express their genes differently, leading to distinct cellular outcomes. Cellular differentiation is the biological process that leads a cell to opt for a particular cellular identity [1]. In contemporary biology, the cellular identity is defined by the expression of cell markers, mostly surface proteins.

Recently, single-cell RNA sequencing (scRNA-seq) has enabled the simultaneous measurement of gene expression levels for up to millions of individual cells and tens of thousand of genes [2, 3, 4]. Thanks to this technique, a cell can be represented in a high-dimensional space, where each dimension corresponds to the expression level of a given gene, so the space dimension equals the number of genes. Repeating such measurements at different time points gives access to the temporal variation, or transport, of a distribution on a gene expression space. The entire temporal trajectory of the distribution characterizes the differentiation process at population level, but trajectories of individual cells are still out of reach since measurement process is destructive [5].

In practice, gene expression is measured either as the number of mRNA molecules or the level of protein expression. Hence, the above-mentioned gene space is either a mRNA counts space or a protein level space. While the differentiation process is fundamentally a continuous process, most of the experimental measurements provide access only to discrete temporal distributions. In order to study a differentiation process and more specifically to infer differentiation pathways, many works focused on the reconstruction of the continuous transcriptomics evolution based on such distributions.

Schiebinger et al. [5] were pioneers in using the optimal transport (OT) theory to infer such trajectories from time-stamped transcriptomics data, introduced in the Waddington Optimal Transport (WOT) algorithm. WOT enables the reconstruction of a “timeline” between two cell distributions measured at different times. WOT solves the OT problem in a standard way: by minimizing a cost function defined as the squared Euclidean distance between cells in a PCA-reduced space.

Since then, several works proposed methods of cell trajectory inference inspired by OT theory [6, 7, 8, 9, 10, 11, 12, 13, 14, 15, 16, 17]. Authors of OT-based studies chose to work with quadratic OT, which consists in using the squared euclidean distance as a cost in the OT problem (see Section 2.3). More specifically, they used the socalled entropic approximation of OT – which is equivalent to solving the Schrödinger problem – in order to use the efficient Sinkhorn algorithm [18, 19]. The Schrödinger problem consists in finding the stochastic process that is the closest to a given reference process while fitting the observations. The reference process is in general a Brownian motion, hence it is implicitely assumed that cell trajectories are close to diffusion. In some cases, distances different from the square Euclidean distance have been considered, mostly for taking into account additional information like lineage information [11, 20] or coexpression between genes [21].

Some methods nevertheless used a different cost function when solving the OT problem. For instance, Liu et al. [17] criticized the use of Euclidean distance when applying OT theory and proposed an implementation of a learned cost function in the OT problem based on the Sinkhorn algorithm [18]. However, they do not use OT theory to infer cell trajectories, instead they focus on alignment of scRNA-seq datasets. Additionally, the learned cost is non-interpretable, a property that is shared by computational methods based on machine learning. In order to reduce the computational cost and gain interpretability, scEGOT [13] combines OT with cell state graph. By adding Gaussian mixture distribution on each cell’s dots, scEGOT allows to consider cell clusters instead of individual cells. The Gaussian mixture makes the results interpretable, however their objective are clearly different from ours because the authors use Euclidean distance between Gaussian distributions to infer trajectories, relying once again on a standard approach.

The main idea of the present paper is to assert that in order to reconstruct cell differentiation trajectories, a more realistic, quantitative model of stochastic gene expression should be considered as reference process. We focus on solving the Schrödinger problem while using an alternative reference process that would be closer to realistic gene dynamics. Indeed, the Diffusive process generates unrealistic dynamics for proteins and mRNA [22]. The Schrödinger problem is not seen here as an approximation of a transport problem, but as a way to identify a process close to an *a priori* quantitative model that would fit the observations.

When examined at the single-cell level, gene expression is undoubtedly a stochastic process [23, 24, 25, 26, 27, 28]. The simplest model that can correctly reproduce experimental data distribution is the well-known two-state model of gene expression [29, 30, 31], a refinement of the model introduced in 1991 by Ko [32]. In this model, a gene is described by its promoter which can be either active or inactive – possibly representing a transcription complex being “bound” or “unbound” although it may be more complicated [33] – with mRNA being transcribed only during the active periods.

Genes may mechanistically influence the activation rate of other genes, thus creating a Gene Regulatory Network (GRN). We consider an extension of the two-state model for several genes, that takes into account the GRN structure. For certain parameter values, mRNA dynamics are characterized by brief and important production of RNA, with long periods of inactivity in between: this regime corresponds to the widely observed phenomenon called transcriptional bursting [29]. Herbach et al. [34] introduced a mathematical framework for modeling GRN driven by transcriptional bursting, which we call the Bursty model here. This model has been shown to reproduce realistic gene dynamics at the single-cell level [35, 36] and will provide our reference process.

In this work we compare the performances of a Diffusive process, associated to standard OT theory, and a reference process based on the Bursty model to solve the Schrödinger problem, when initial and final distributions are based on realistic, simulated data. We first explore the case of a single gene, then of a specific two-gene GRN, the toggle switch, and finally more complex three-gene GRNs. We focus on the influence of some indicators, namely the time difference between the measurements of the two distributions and the number of cells in the distributions. We show that using the Bursty model as a reference process provides a better solution to the Schrödinger problem for all tested networks.

## 2 Methods

### 2.1 Gene dynamics

We model gene expression using the framework introduced by Herbach et al. [34], and later simplified in the bursty regime [37, 38, 39]. In this model, a gene is characterized by its amount of mRNA and its level of proteins: mRNA is synthesized through stochastic transcriptional bursts, and degraded at rate *d*_0_; proteins are synthesized from mRNA at rate *s*_1_ and degraded at rate *d*_1_. The Bursty model is a piecewise-deterministic Markov process, with deterministic part given by

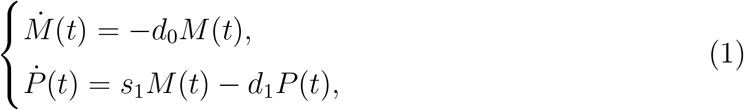

where *M*(*t*) is the quantity of mRNA and *P*(*t*) the quantity of protein at time *t*, and a stochastic part: mRNA bursts occur at random times with rate *k*_*on*_ and have a random size that follows an exponential distribution [38, 39].

Within a cell, genes interact with each other via a GRN. Hence, a gene can be activated or inhibited by itself but also by other genes in the GRN. A burst is assumed to occur for gene *i* at a rate *k*_*on,i*_(*P*_1_, …, *P*_*G*_), where *G* is the number of genes in the GRN and (*P*_1_, …, *P*_*G*_) is the vector of protein levels of all genes in the GRN. The burst rate of gene *i* accounts for a basal activity parameter *β*_*i*_ and the influence of other genes in the GRN, expressed via a gene-gene interaction matrix *θ* = {*θ*_*ji*_}_*j,i*∈{1,…, *G* }_ [38, 40], so that

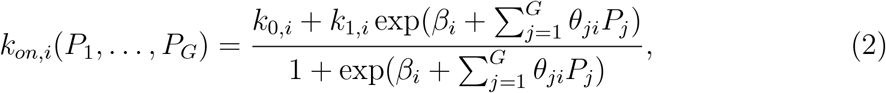

where *k*_0, *i*_ and *k*_1, *i*_ correspond respectively to the minimum and maximum burst frequencies of gene *i*, and *θ*_*ji*_ is the intensity of gene *j* action on gene *I* [34]. Parameters *θ*_*ji*_ can be positive or negative, depending on whether gene *j* activates or inhibits gene *i*(Figure 1).

**Figure 1:**
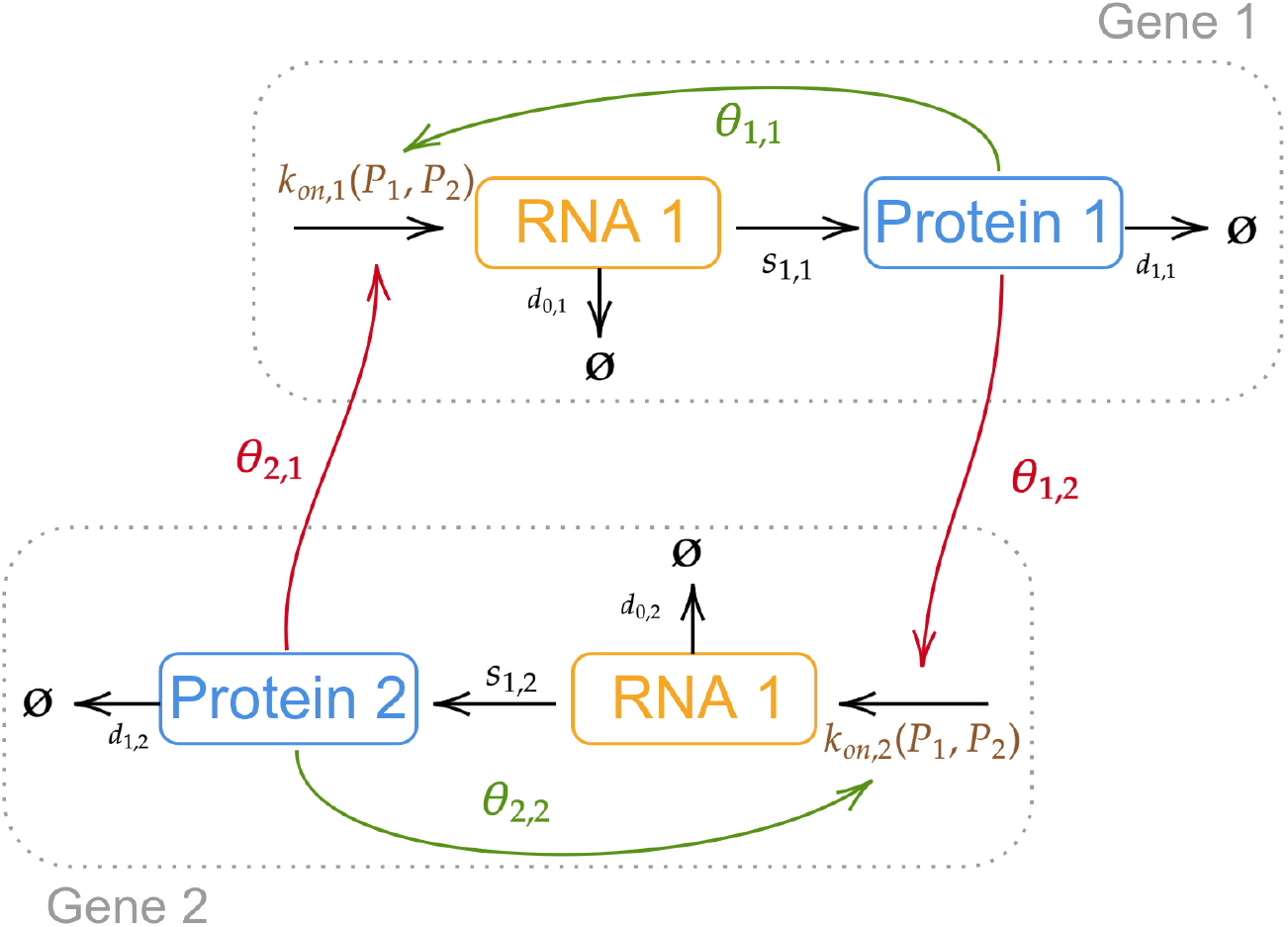
Schematic representation of a two-gene GRN based on the Bursty model of gene dynamics. For each gene *i* = 1,2, mRNA synthesis is activated with a rate *k*_*on,i*_(*P*_1_, *P*_2_), defined in (2), and mRNA and protein dynamics follow system (1). Parameters of the GRN are described in the text. In the case of the toggle switch model, the GRN consists of 2 genes that self-activate (*θ*_1,1_ > 0 and *θ*_2,2_ > 0, green arrows) and inhibit each other (*θ*_2,1_ < 0 and *θ*_1,2_ < 0, red arrows).

#### Example of GRN: the Toggle Switch

We consider a standard GRN made of two genes, the so-called “toggle switch” model. The two genes self-activate and inhibit each other, i.e.

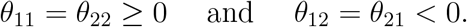

This GRN is illustrated in Figure 1 and Figure 2 illustrates the temporal evolutions of mRNA and protein for a toggle switch GRN where gene dynamics follow the Bursty model, computed with parameters values in Table 1. A deterministic toggle switch model would favor one gene over the other, depending on initial conditions of the system. In this stochastic version of the toggle switch, one observes that one gene activity always prevails, yet the dominant gene switches stochastically.

**Table 1:** Default parameter values.

| Description | Parameter | Value | Unit |
| --- | --- | --- | --- |
| Number of genes | $G$ | 2 | genes |
| Number of cells | $N$ | 100 | cells |
| Number of simulations | $n$ | 100 | simulations |
| Initial conditions for mRNA levels | $IC_M$ | (0, 0) | mRNA |
| Initial conditions for protein levels | $IC_P$ | (0, 0) | protein |
| Smoothing standard deviation | $\varepsilon$ | $10^{-4}$ | protein |
| Creation rate for proteins of gene $i$ | $s_i$ | 10 | protein h<sup>-1</sup> mRNA<sup>-1</sup> |
| Minimum burst frequency of gene $i$ | $k_{0,i}$ | 0 | h<sup>-1</sup> |
| Maximum burst frequency of gene $i$ | $k_{1,i}$ | 2 | h<sup>-1</sup> |
| Basal activity of gene $i$ | $\beta_i$ | 5 | h<sup>-1</sup> |
| Degradation rate of mRNA of gene $i$ | $d_{0,i}$ | 10 | h<sup>-1</sup> |
| Degradation rate of protein of gene $i$ | $d_{1,i}$ | 5 | h<sup>-1</sup> |
| Intensity of self-activation of gene $i$ | $\theta_{ii}$ | 10 | |
| Intensity of inhibition of gene $i$ | $\theta_{ij}$ | -10 | |
| Initial time point in transient state | $t_1$ | 0.1 | h |
| Final time point in transient state | $t_2$ | 0.5 | h |
| Diffusion coefficient | $\sigma^2$ | 0.001/2 | protein<sup>2</sup> h<sup>-1</sup> |
| Precision of Sinkhorn algorithm |  | 0.1 |  |

**Figure 2:**
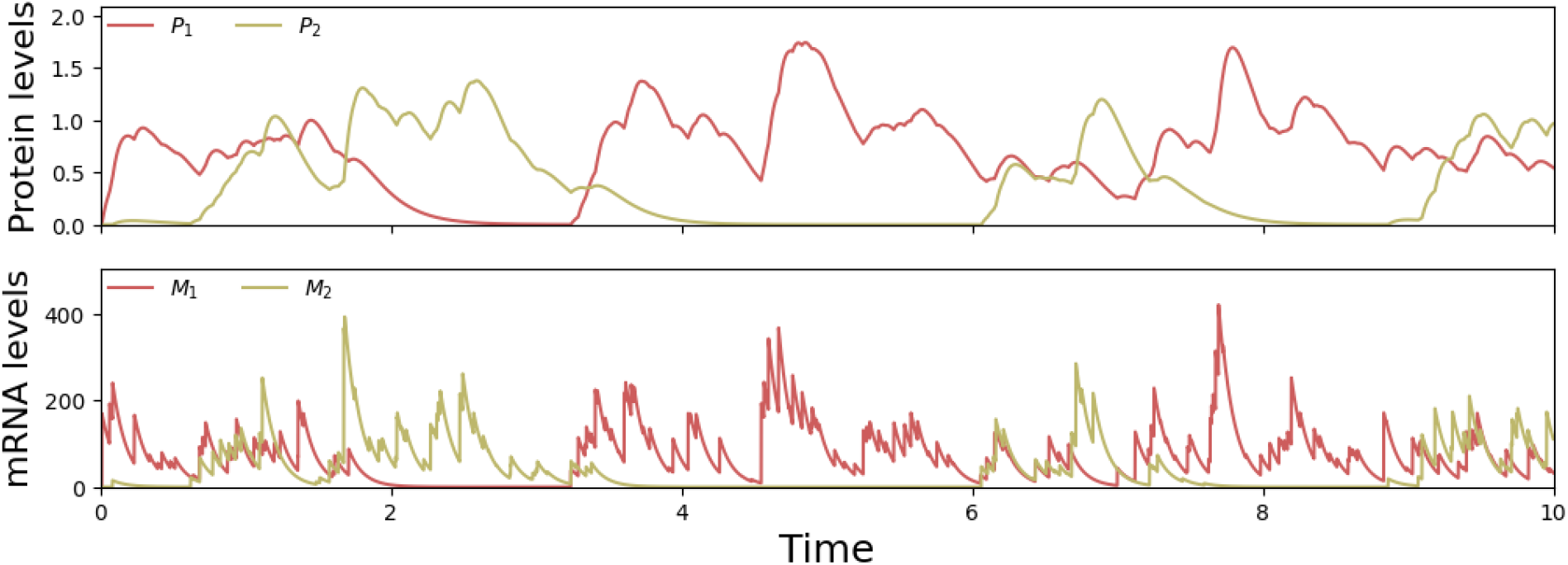
Temporal evolution of protein and mRNA levels for the toggle switch model. Dynamics of one gene are illustrated in yellow, the other in red. Protein levels are displayed on the upper panel, while mRNA dynamics are displayed in the lower panel

More generally, by modifying the values of the interaction parameters *θ*_*ji*_ in model (1)-(2), any two-gene GRN can be implemented (see, for example, Figure 10). Extensions to three-gene GRNs can be straightforwardly implemented (see, for example, Figure 11).

#### Simulations of protein dynamics

Computations of the temporal evolution of protein and mRNA levels were performed using the Python package Harissa [40], which implements an exact simulation procedure for the Bursty model [37] (Figure 2). This allows to simulate one GRN, hence simulating *N* times the evolution of a GRN results in *N* independent cell trajectories (Figure 3).

**Figure 3:**
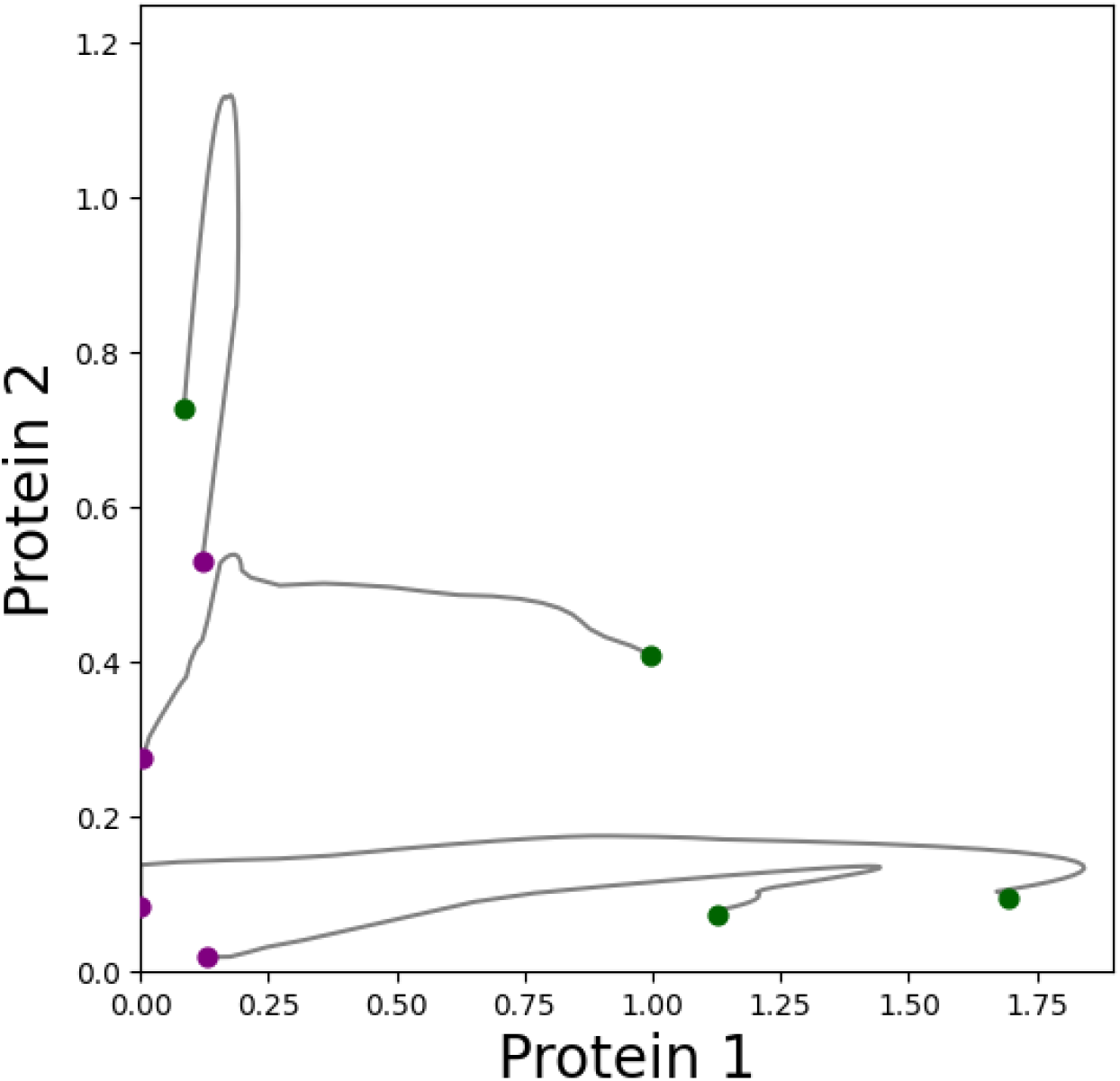
Cell trajectories of four cells driven by a toggle switch. Initial (time *t*_1_ = 0.1h) location of cells is displayed as purple dots, while their final (time *t*_2_ = 0.5h) location is represented by green dots. Each grey line is the temporal trajectory of each cell. Parameters are given in Table 1.

In order to mimic biological experiments aimed at measuring a GRN activity at given observation times, we simulate *N* times the GRN with initial conditions *IC*_*M*_. and *IC*_*P*_ (see Table 1), select a simulation time *t*, and extract all mRNA and proteins concentrations for all genes.

Throughout this work, we focus on distributions of protein levels only, yet the case of mRNA distributions is discussed in the last section. We denote by *µ* the distribution of *G* protein levels in a population of *N* cells at time *t* = *t*_1_, and *ν* the distribution of *G* protein levels in the same population at time *t* = *t*_2_, with *t*_2_ > *t*_1_. They are given by:

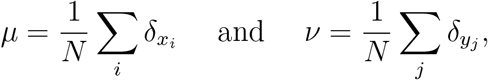

whereδis the Dirac distribution,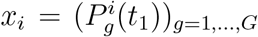_, *G*_ is the level of gene *g*’s protein in cell *x*_*i*_ at *t*_1_ and 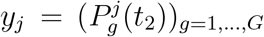, is the level of gene *g*’s protein in cell *y*_*j*_ at *t*_2_. For the sake of simplicity, cells have the same index in distributions *µ* and *ν*(i.e., the *i*-th component of *µ* is associated to the same cell than the *i*-th component in *ν*). Additionally, the two distributions are jointly normalized (between 0 and 1). Default parameter values are given in Table 1.

### 2.2 Reduced model with analytical solution

In addition to the model introduced in the previous section, for which couplings will be estimated from simulations, we consider a simplified model (*reduced Bursty model*) that will provide closed-form couplings. This simplification is achieved in two steps. First, a protein-only model is derived from a time scale separation of mRNA and proteins corresponding to *d*_0_ ≫ *d*_1_ (i.e., proteins assumed to be more stable than mRNAs). This model is defined by [40, 41]:

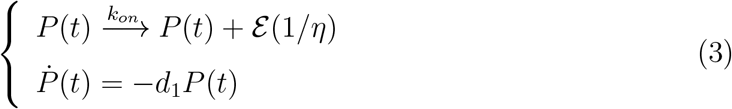

Where ℰ is the exponential distribution characterizing the random size of each burst (with mean value *η*), *k*_*on*_ is the burst rate and *d*_1_ the degradation rate of proteins, as before. Second, we consider a particular 1-dimensional case of model (3) with a constant *k*_*on*_: this setting can be interpreted either as a single gene with no feedback, or part of a GRN within a time range sufficiently short so that *k*_*on*_(*P*_1_, …, *P*_*G*_) does not vary significantly during this period.

This model turns out to be analytically tractable (see Supplementary File). The time-dependent distribution of *P*(*t*) knowing *P*(0) = *P*_0_ is given by

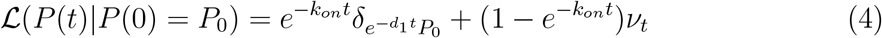

Where 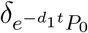 is the Dirac measure localized at 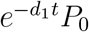, and *ν*_*t*_ has a density function *g*_*t*_(*x* | *P*_0_) given by

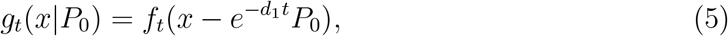

where

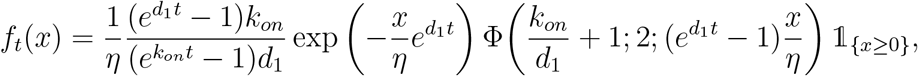

and Φ denotes the confluent hypergeometric (Kummer’s) function of the first kind.

For numerical applications, we regularize the singular part in (4) by replacing it with a normal distribution with small standard deviation *ε* ≪ 1,

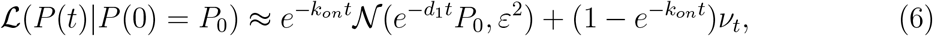

so that the equality holds when *ε* → 0.

### 2.3 Coupling between time points

Let’s recall that experimental data would be collections of cell positions in the protein space at distinct time points, and that the objective of cell trajectory inference is to reconstruct cell trajectories in this space. Hence, we restrict the problem to two consecutive time points *t*_1_ and *t*_2_. Given two collections 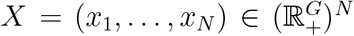 and 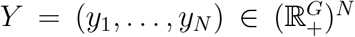, we search to construct a matrix 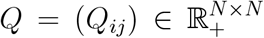 of probabilities *Q*_*ij*_ that a cell at position *x*_*i*_ at time *t*_1_ reaches the state *y*_*j*_ at time *t*_2_, in the most realistic way. In the following, the matrix *Q* will be called a coupling between the collections *X* and *Y*.

By solving the Schrödinger problem (see Section 2.4), we look for the coupling that is the closest to a certain reference coupling 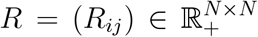, obtained using the Bursty model (1), while being compatible with the data *X* and *Y*. For either the model (1) or the Diffusive model, the reference coupling *R* is built by associating to the model a function being an exact or an approximated version of the transition kernel of the underlying theoretical process.

#### Coupling for the reduced Bursty model

In the case of a single gene following model (3), we use the regularized analytical solution (6) to compute the probability density of reaching state *y* starting from *x*, that is *p*_*t*_(*y* | *x*) defined by

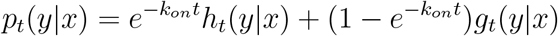

where *g*_*t*_ is defined in (5) and

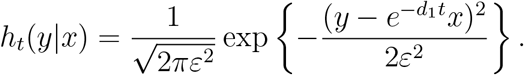

Considering this transition kernel *p*_*t*_(*y* | *x*) and the uniform distribution *p*(*x*) ≡ 1 */N* over initial states (*x*_1_, …, *x*_*N*_), the related reference coupling *RB*_ref_(*x, y*), for “reduced Bursty reference process”, is defined as:

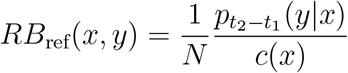

where the normalizing term

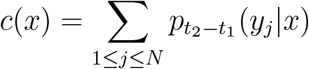

is defined so that 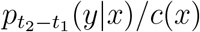 is a discrete probability distribution over (*y*_1_, …, *y*_*N*_).

#### Coupling for the non-reduced Bursty model

Given a GRN whose dynamics are given by a Bursty model (1)-(2), we define *B*_ref_(*x, y*) an approximation of its coupling between two collections at two different time points. It is the reference process defined by the probability of reaching *y* when starting from *x*, computed from the Bursty model (1)-(2).

We estimate *B*_ref_ using a Gaussian method (illustrated in Figure 4). For each cell *i* in the initial distribution *µ* at *t* = *t*_1_, we simulate the Bursty model (1)-(2) *n* times with the cell *i*’s coordinates as an initial condition. We obtain a distribution denoted 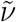 at time *t* = *t*_2_, representing the potential fates of cell *i*. An estimated density of 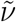 is computed using Gaussian approximation, i.e. we create Gaussian distributions centered on cells in the distribution 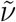, and we compute the Gaussian kernel of this distribution using the default parameters of the textitgaussian kde() python function. The result is then divided by the total sum to ensure the resulting distribution is a probability distribution.

**Figure 4:**
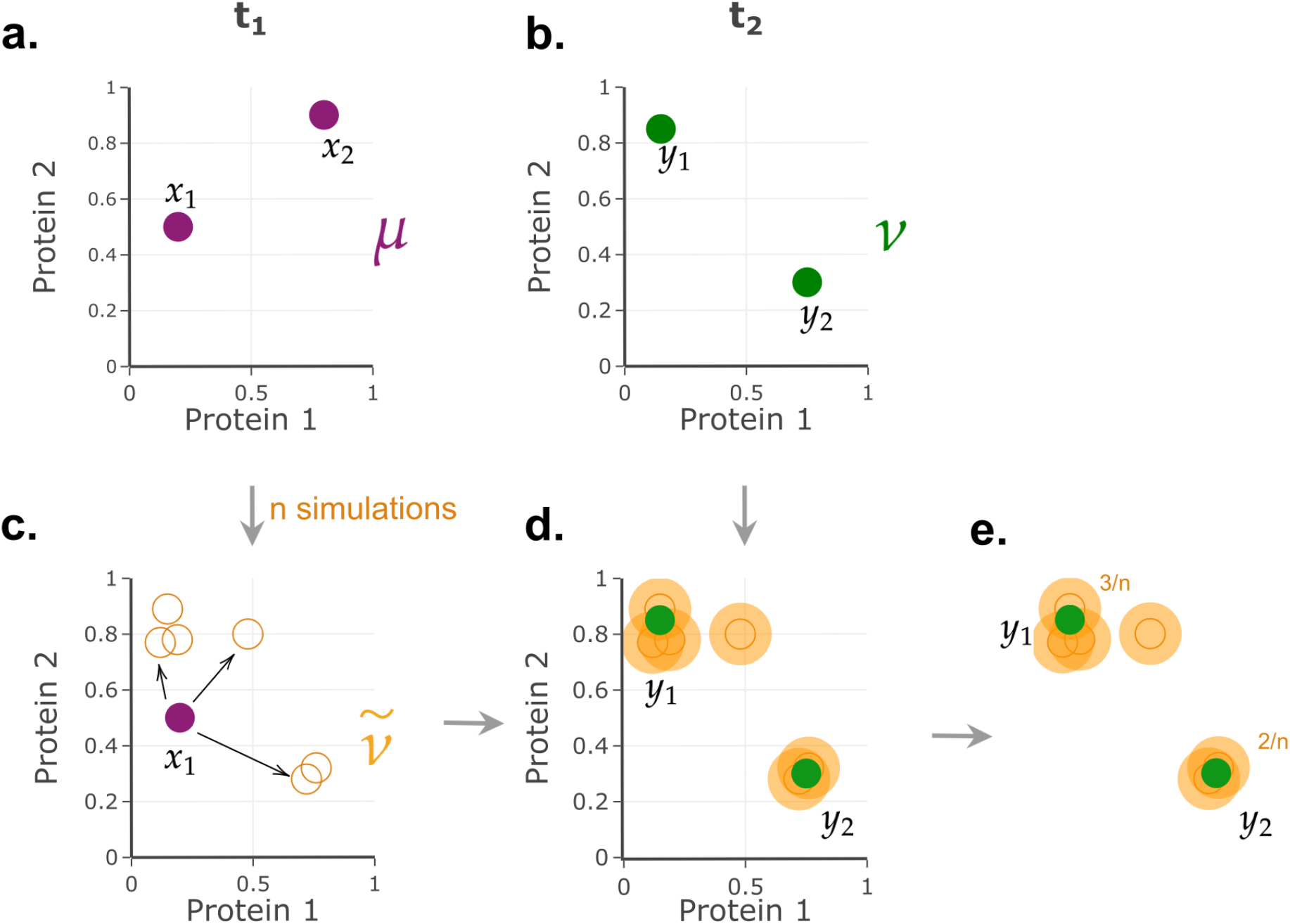
Schematic of the procedure used for constructing the *B*_ref_ estimation with a Gaussian method. **a**. Cells *x*_1_ and *x*_2_ (purple dots) are two cells of the distribution *µ* at time *t*_1_ in protein space. **b**. Cells *y*_1_ and *y*_2_ (green dots) are two cells belonging to the distribution *ν* at time *t*_2_ in the protein space. **c**. The approximated distribution 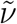 is computed for *n* = 6 and cells are displayed as orange circles. **d**. Gaussian distributions centered on 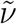 cells generate a smooth distribution (orange areas). **e**. Probabilities *p*(*y*_1_ *x*_1_) = 3 */n* and *p*(*y*_2_ *x*_1_) = 2 */n* are deduced from the construction process.

#### Coupling for a Diffusive process

When the underlying model is a Diffusive process, we consider the natural discrete counterpart used in [6], that is the reference coupling denoted by *D*_ref_ is defined by

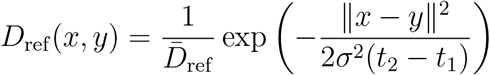

where *σ* ^2^ is the diffusion coefficient,∥ *x* − *y* ∥is the Euclidean distance (here, in protein space) between *x* and *y*, and 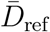 is the normalizing constant given by

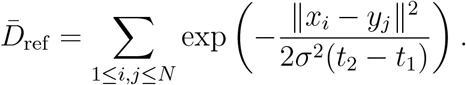

The value of the diffusion coefficient was chosen to be both relevant and close to the value used in Schiebinger et al. [5]. The challenge was to be small enough for the convergence of the Schrödinger problem while using a diffusive process that still predicts couplings (*D*_ref_(*x, y*)> 0).

### 2.4 Schrödinger problem

Introduced in 1932 [42], the Schrödinger problem is an entropic minimization between a stochastic process and a reference process with marginal constraints. This problem has been studied because when the reference process is a Brownian and the diffusion coefficient goes to 0, it converges to the quadratic OT problem [43, 44].

Let Π and Ψ be two discrete and finite spaces, e.g. Π = { *x*_1_, …, *x*_*N*_ } and Ψ = { *y*_1_, …, *y*_*N*_ } with *N* ∈ℕ, the number of cells. We denote by *R* ∈ 𝒫 (Π×Ψ), the reference process, where 𝒫 (Π×Ψ) is the space of probability measures on Π×Ψ. Since Π and Ψ are discrete spaces of size *N*, then *R* is a *N* × *N* matrix. We denote by *u* ∈ 𝒫 (Π) and *v* ∈ 𝒫 (Ψ) two uniform vectors of size *N*(each cell has the same weight), and we search for the process *Q* solution of

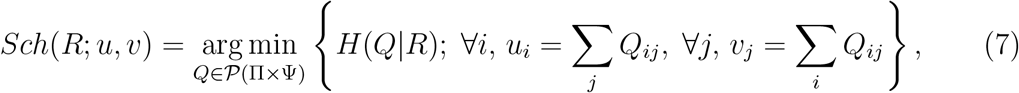

where *H* is the *relative entropy* of *Q* with respect to *R*(also called Kullback–Leibler divergence of *Q* from *R*) defined by

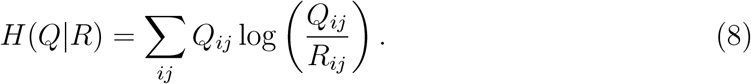

This quantity will be called *entropy* throughout this paper for simplicity. The Schrodinger problem (7) can be solved with the Sinkhorn algorithm [18, 19].

We solve the Schrödinger problem for the reference processes introduced in Section 2.3. When *R* = *RB*_ref_,

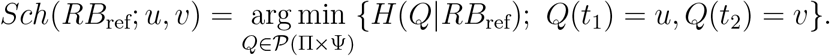

We denote by *RB*_sch_ its solution, *RB*_sch_:= *Sch*(*RB*_ref_; *u, v*).

For the reference processes *R* = *B*_ref_, we write

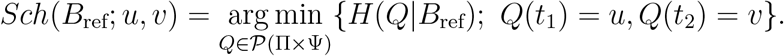

We denote by *B*_sch_ its solution, *B*_sch_:= *Sch*(*B*_ref_; *u, v*).

When *R* = *D*_ref_, then

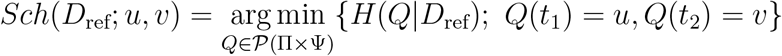

and we denote *D*_sch_ its solution, *D*_sch_ = *Sch*(*D*_ref_; *u, v*).

### 2.5 Performances of the models

In order to compare the performances of the Bursty model with other models, in particular the Diffusive model of gene dynamics, we rely on the computation of a standard criterion, the entropy (8) between solutions of the Schrödinger problem (7). However, to ensure that results do not depend on the criterion, we also test other criteria measuring the difference between two stochastic processes.

#### Entropy between Schrödinger problem solutions

For one-gene GRN, we define

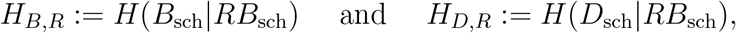

i.e. the entropy between *B*_sch_ or *D*_sch_ and *RB*_sch_. This value characterizes the difference between the ability of a Diffusive process or a Bursty model-based process versus *RB*_sch_ to predict cell trajectories correctly. The process *RB*_sch_ is considered as the reference process because it provides an approximation of the exact probability distribution. In parallel, we compute as a control measure

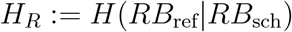

in order to estimate the error made when performing stochastic simulations.

For two-or-more-gene GRN, we define

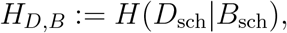

the entropy between *D*_sch_ and *B*_sch_, in order to evaluate the difference between the ability of a Diffusive process versus a Bursty model-based process to correctly predict cell trajectories. Similarly to the one-gene GRN case above, we introduce

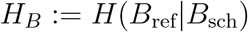

to estimate the error made when performing stochastic simulations.

Values of *H*_*B*_ should be smaller than *H*_*D,B*_ values if *B*_ref_ and *B*_sch_ are closer than *D*_sch_ and *B*_sch_. If the Bursty model-based reference coupling better predicts individual cell trajectories, the difference between *H*_*B*_ and *H*_*D,B*_ will be important.

Due to stochasticity, we perform 100 iterations to compute these quantities and focus on the mean and standard deviation.

#### Other difference indicators

To certify that our results do not depend on the choice of the entropy, we also compute the following criteria, for two processes *Q* and *R*:

- the Total Variation Distance:

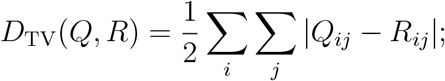
- the Jensen–Shannon Divergence:

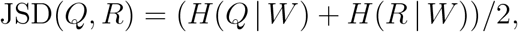

where *W* = (*Q* + *R*) */* 2;
- the Frobenius norm:

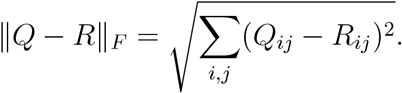

## 3 Results

We consider two distributions of protein levels obtained from a fixed number of cells, simulated from the Bursty model (see Section 2.1) and measured respectively at times *t*_1_ and *t*_2_ (*t*_2_ > *t*_1_). Noticeably, we focus only on protein levels since protein dynamics, contrary to mRNA dynamics, allow to preserve the markovian property in the Bursty model therefore providing a rigorous framework. We solve the associated Schrödinger problem for several reference processes, including the Bursty process and the standard Diffusive process. Solutions of the Schrödinger problem are compared using an entropy function. If the two processes are similarly able to correctly infer trajectories, the entropy equals or nears zero.

When the final distribution is measured a “long” time after the initial distribution, the Bursty model has reached a so-called stationary state, and both initial and final distributions can be considered independent. In this case, trajectory inference should depend less on the selected reference process. Consequently, we focused our study on distributions in transient states, meaning the time difference *t*_2_ − *t*_1_ is small enough and *t*_1_ is close to the initialisation of the simulations, as represented in Figure 3. Details are provided in Section 2.

### 3.1 Schrödinger problem for a single gene

We first considered the case of a single gene with no feedback, as a toy model to illustrate the influence of the underlying model of gene dynamics on trajectory inference.

Figure 5 shows the solutions of the Schrödinger problem computed with different approximated processes: a Diffusive model, *D*_sch_ (Figure 5a), the non-reduced Bursty model, *B*_sch_ (Figure 5b), and the reduced Bursty model, *RB*_sch_ (Figure 5c). The ground truth (Figure 5d) is a coupling of cells with the same index (cell *i* in the initial distribution is cell *i* in the final distribution). We observe that *RB*_sch_ yields the closest result to the ground truth (Figure 5c), in the form of a very accurate transition matrix, while *D*_sch_ yields the furthest (Figure 5a). A darker diagonal is observed for *B*_sch_ (Figure 5b), meaning that this process better approximates the ground truth than *D*_sch_. Noticeably, a limitation of the representation in Figure 5 is that the coupling could be not expressed (light color) and yet be relevant. For example, if two cells are close in protein space at both *t*_1_ and *t*_2_, then the coupling between those cells could be mixed up but relevant.

**Figure 5:**
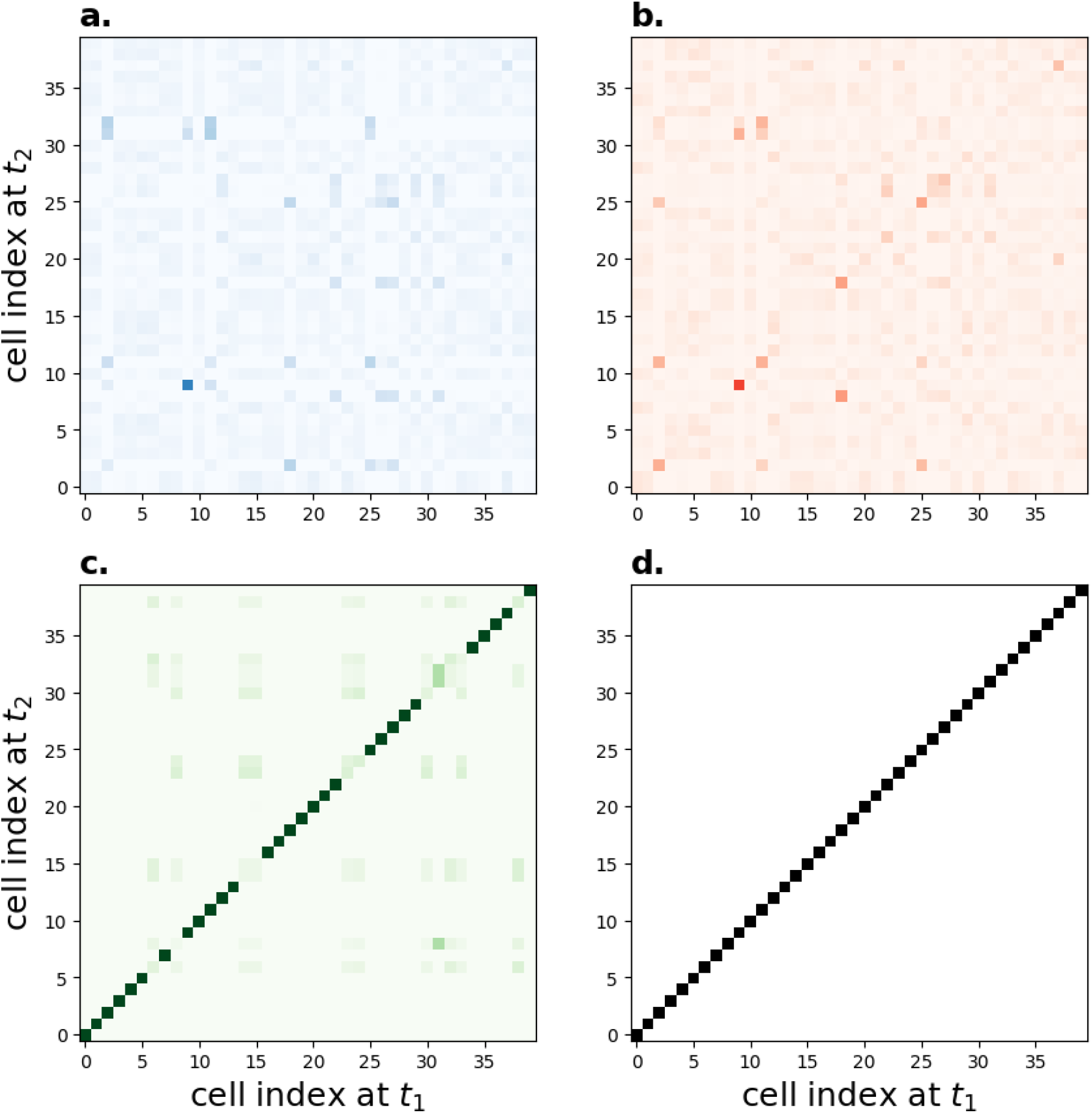
Solution of the Schrödinger problem for a single gene. **a**. *D*_sch_. **b**. *B*_sch_. **c**. *RB*_sch_.**d**. Ground truth. Each colored square represents the probability of the cell index at *t*_2_ starting from the cell index at *t*_1_ for *N* = 40 cells: the darker the box, the greater the probability. All matrices sum to 1. Parameters are given in Table 1.

Figure 6 presents the entropy between *D*_sch_ and *RB*_sch_, denoted by *H*_*D, R*_, and the entropy between *B*_sch_ and *RB*_sch_, denoted by *H*_*B, R*_ (see Section 2.5). The curve *H*_*R*_ also appears in Figure 6, it serves as a control by computing the entropy between the reference process and its approximation. The initial distribution is measured at time *t*_1_ = 0.1 h, and the final distribution is computed at various times *t*_2_ in the range [0.2; 0.5] h.

**Figure 6:**
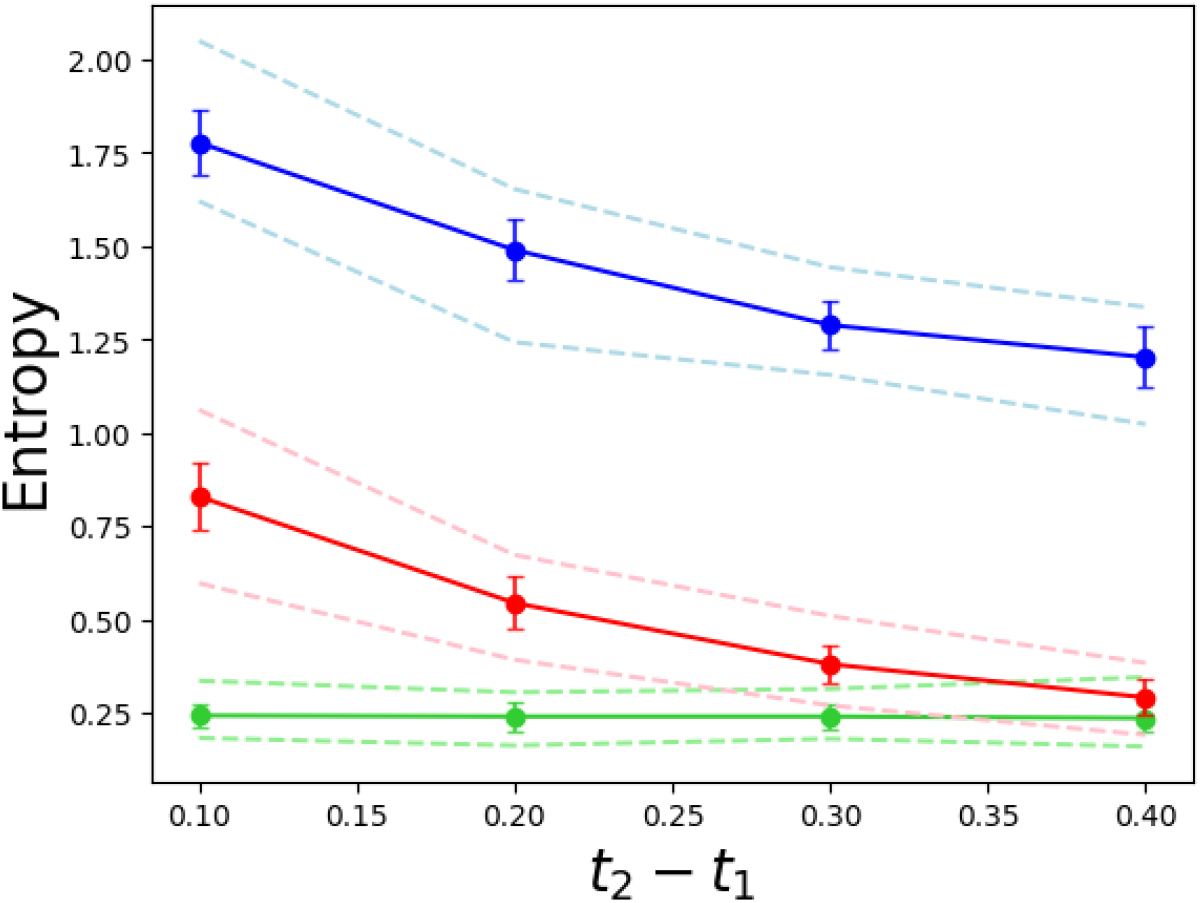
Entropies between reference processes as a function of the time interval for a one-gene GRN. Mean and standard deviation (straight lines) of entropies *H*_*D, R*_ (blue), *H*_*B, R*_ (red) and *H*_*R*_ (green) are displayed for various values of the time difference *t*_2_ − *t*_1_, when the gene dynamics model is in transient state. Additionally, minimum and maximum values over 50 iterations (dashed lines) are also displayed. Other parameters are detailed in Table 1.

We observe that quantities *H*_*D, R*_ and *H*_*B, R*_ decrease as *t*_2_ − *t*_1_ increases and *H*_*R*_ remains constant. We also observe that *H*_*D, R*_ is always higher than the entropy between *B*_sch_ and *RB*_sch_, and remains much higher than *H*_*R*_ even for large values of *t*_2_ − *t*_1_ (here 0.4 h). Entropy *H*_*B, R*_ is always larger than *H*_*R*_, yet it is much smaller than *H*_*D, R*_, and for *t*_2_ − *t*_1_ = 0.4 h there is no difference between *H*_*B, R*_ and *H*_*R*_. Both the exact temporal distribution of the process and its approximation provide comparable results.

The study of this toy model highlights that using the exact temporal distribution of the reference process to solve the Schrödinger problem provides much better results than using the approximation of a gene dynamics model. However, it is a complex, most of the time impossible, task to find the analytical expression of this distribution for a GRN comprising at least two connected genes. Consequently, for a more relevant GRN, we will focus on the estimations provided by the reference process of the non-reduced Bursty model and the Diffusive process.

### 3.2 Case study: the toggle switch

We focused on a two-gene GRN, called ‘toggle switch’, introduced in Section 2.1 and illustrated in Figure 10b (blue GRN). This specific GRN consists of two self-activating genes that inhibit each other.

#### 3.2.1 Influence of the time interval and the number of cells

We first investigated the influence of the time between measurements of the initial and the final distributions (i.e. *t*_2_ − *t*_1_) on the ability of two models, the Bursty and the Diffusive models, to correctly transport cell distribution.

Figure 7a shows values of entropies *H*_*D,B*_ and *H*_*B*_ as functions of the time difference *t*_2_ − *t*_1_. The *H*_*D,B*_ curve is always above the *H*_*B*_ curve and remains approximately constant and independent of the time difference. The entropy between the two processes is the same for all *t*_2_ − *t*_1_ values: whatever the time difference, the Diffusive process infers trajectories with the same uncertainty. For the time difference *t*_2_ − *t*_1_ = 0.2h, the entropy *H*_*D,B*_ has a larger standard deviation, due to one extreme value.

**Figure 7:**
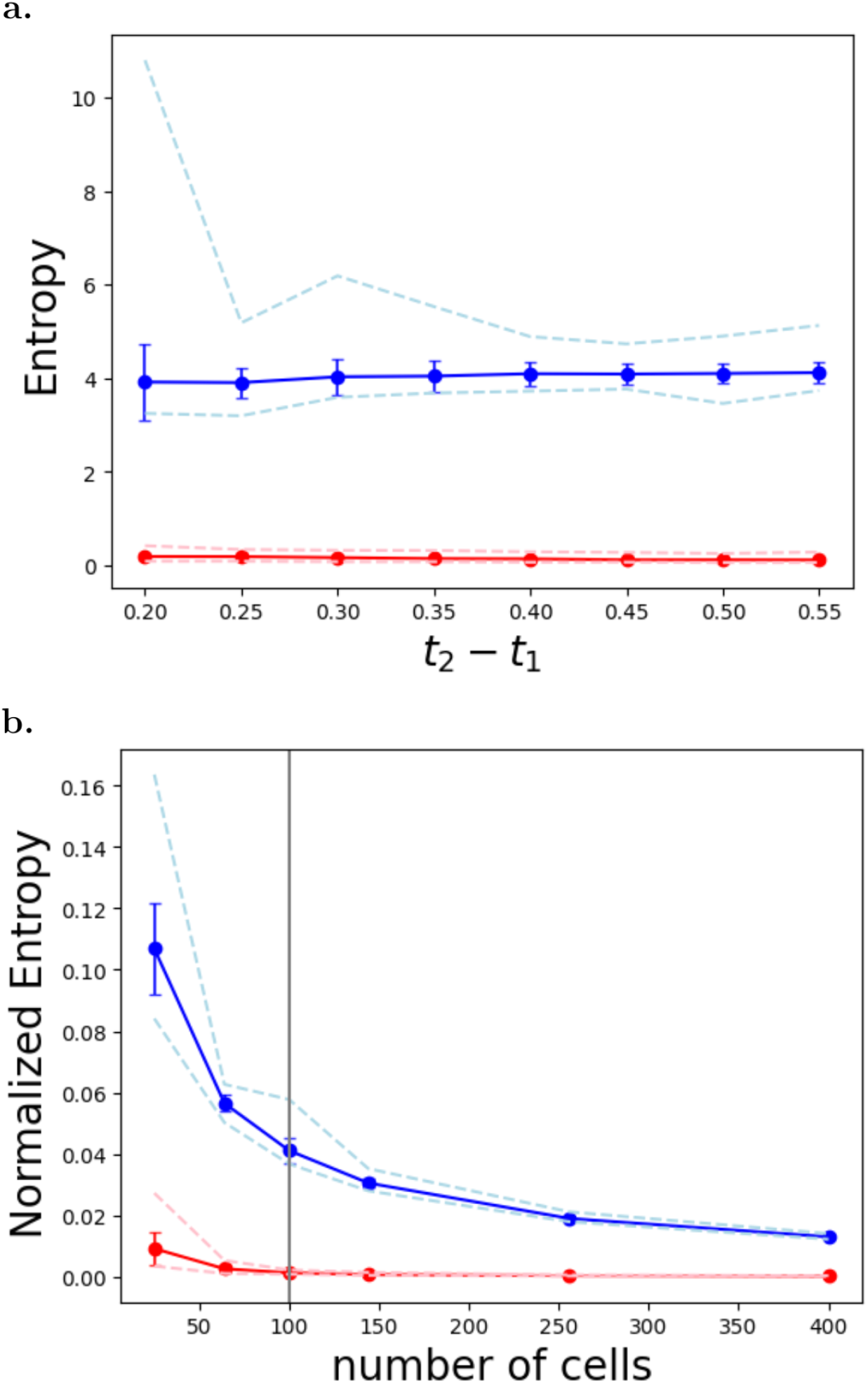
Entropies between reference processes for the toggle switch GRN, in the transient state. Mean and standard deviation (straight lines) of (**a**.) *H*_*D,B*_ (blue) and *H*_*B*_ (red) as functions of the time interval *t*_2_ − *t*_1_, and (**b**.) *H*_*D,B*_ */N*(blue) and *H*_*B*_ */N*(red) as functions of the number of cells *N*. Additionally, minimum and maximum values over 100 iterations are represented (dashed lines). The thick vertical grey line in Panel B is the default value *N* = 100 cells used in other figures. Other parameters are given in Table 1.

Although it is more relevant to focus on the transient state (Figure 7a), it may be informative to question what happens in the stationary state. In this case, cells may either remain in their basin of attraction or switch to the other basin of attraction. The basin of attraction is a notion related to deterministic dynamics, consequently in the current framework where we consider stochastic models of gene dynamics it refers to the basin of attraction of the coarse approximation in the form of a system of ordinary differential equations of a piecewise-deterministic Markov process [45]. Results in the stationary state are illustrated in Supplementary File, they highlight conclusions similar to the ones obtained for the transient state. Although the Diffusive model is less accurate than the Bursty model for inferring trajectories, its ability increases when the final distribution becomes independent of the initial one (later time measurements).

Figure 7b illustrates the influence of the number of cells *N* considered when measuring both initial and final distributions in transient state. To that aim, the entropy has been normalized by the number of cells to avoid a bias due to the size of the collections in the computation of the entropy: the computation of the entropy (8) explicitly depends on the number of cells *N*. First, we observed that *H*_*B*_ */N* does not depend on the number of cells. Second, *H*_*D,B*_ */N* decreases as the number of cells increases. We observed (not shown) that the Diffusive process tends to couple a small sample of cells from the initial collection to all other cells in the final collection. The solution to the Schrödinger problem associated with the reference Diffusive process is then a matrix exhibiting over-coupling for a sample of cells and under- or no-coupling for the rest of the cells. The over-coupled sample is proportionally larger for small numbers of cells, leading to a larger entropy as observed in Figure 7b.

Both numerical investigations on the dependence of *H*_*D,B*_ and *H*_*B*_ on the length of the time interval *t*_2_ − *t*_1_ and the number of cells *N* highlighted a poor ability of the Diffusive model to reconstruct cell trajectories when the underlying model is the Bursty model of gene dynamics. These results are based on the use of the entropy to measure the differences between solutions of the Schrödinger problem. We then questioned the relevance of the entropy criterion and the parameter values of the Bursty model in reaching this conclusion.

#### 3.2.2 Unspecificity of the entropy indicator and the distribution parameters

In Figure 8, we compared the performance of different indicators when evaluating the difference between the Diffusive and the Bursty process. Figure 8 shows mean and standard deviation of normalized indicators, computed as (*Indic* − *ControlIndic*) */ControlIndic*, where *Indic* is either *H*_*D,B*_, *D*_*TV*_ (*D*_*sch*_, *B*_*sch*_), *JSD*(*D*_*sch*_, *B*_*sch*_) or the Frobenius norm ∥ *D*_*sch*_ − *B*_*sch*_∥_*F*_ (see Section 2.5) and *ControlIndic* its associated control measure (respectively, *H*_*B*_, *D*_*TV*_ (*B*_*ref*_, *B*_*sch*_), *JSD*(*B*_*ref*_, *B*_*sch*_) and ∥*B*_*ref*_ − *B*_*sch*_∥_*F*_). We observe that for all indicators, the curves are all strictly positive and increasing. So, despite quantitative differences, all indicators yielded the same qualitative conclusion: the Bursty process provides a better trajectory inference than the Diffusive process.

**Figure 8:**
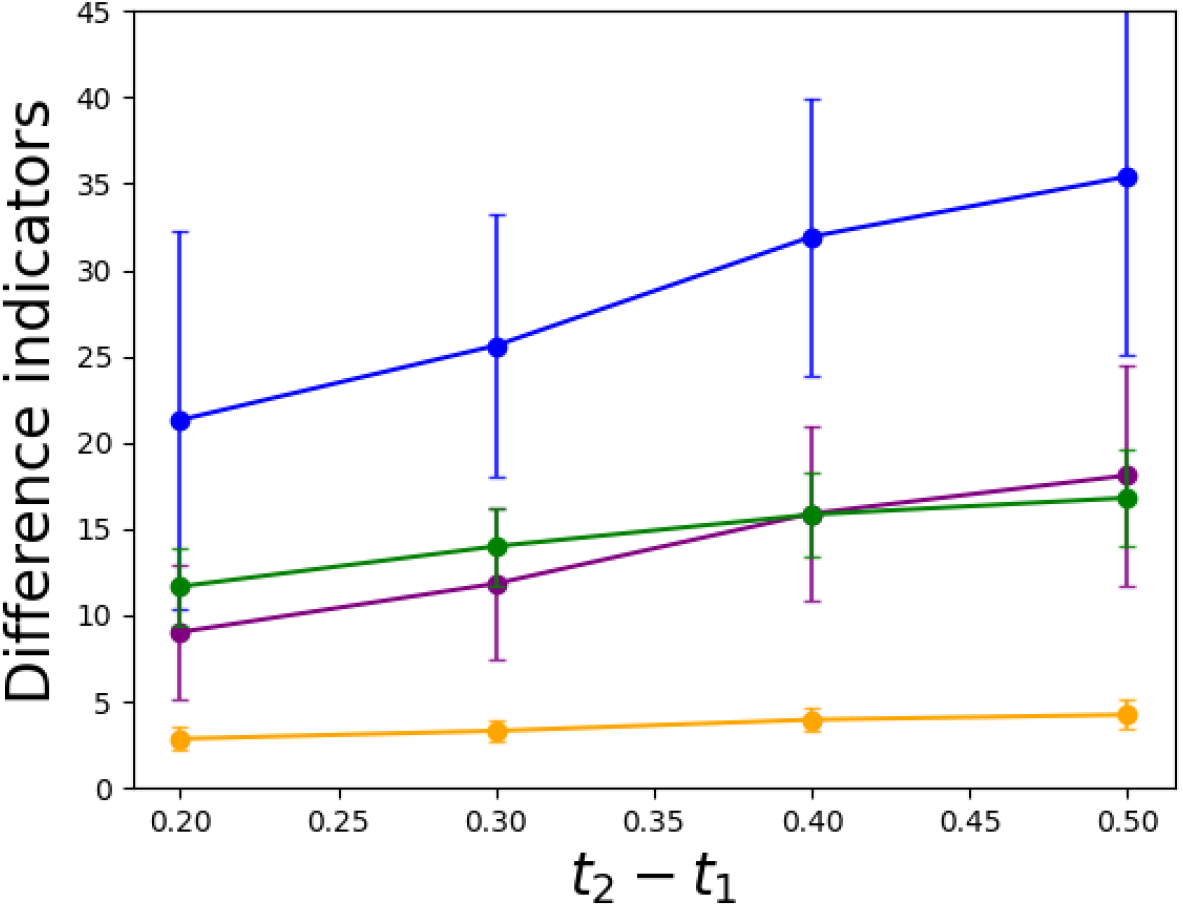
Difference indicators between reference processes as a function of the time interval for the toggle switch GRN. Mean and standard deviation of difference indicators comparing *D*_sch_ and *B*_sch_ normalized by the difference indicators comparing *B*_ref_ and *B*_sch_ respectively: Entropy (blue), total variation distance (yellow), Jensen–Shannon Divergence (purple), and Frobenius norm (green). For standard deviations < 0.1, error bars are not visible. Parameters are detailed in Table 1.

To challenge the performance of the *B*_ref_ process, we tested a different Bursty model as reference process than the one used to generate the two distributions. In practice, we computed the initial distribution *µ* and the final distribution *ν* using the default parameter values of the toggle switch GRN, but we solved the Schrödinger problem with a different *B*_ref_ process (different gene interaction matrix).

Figure 9a highlights that for all the two-gene-based Bursty models (listed in Figure 9c), the curves are very close. The *H*_*D,B*_ curves are all higher than the *H*_*B*_ curves, indicating that any Bursty process, even one with parameter values different from the ones corresponding to the data, provides a better approximation than the Diffusive process. In Figure 9b, one observes that even though *H*_*B*_ curves are nearly indistinguishable at early times they become separated as *t*_2_ − *t*_1_ increases. We observed differences when parameter values of the GRN associated with self-activation were reduced (purple and yellow curves and GRN).

**Figure 9:**
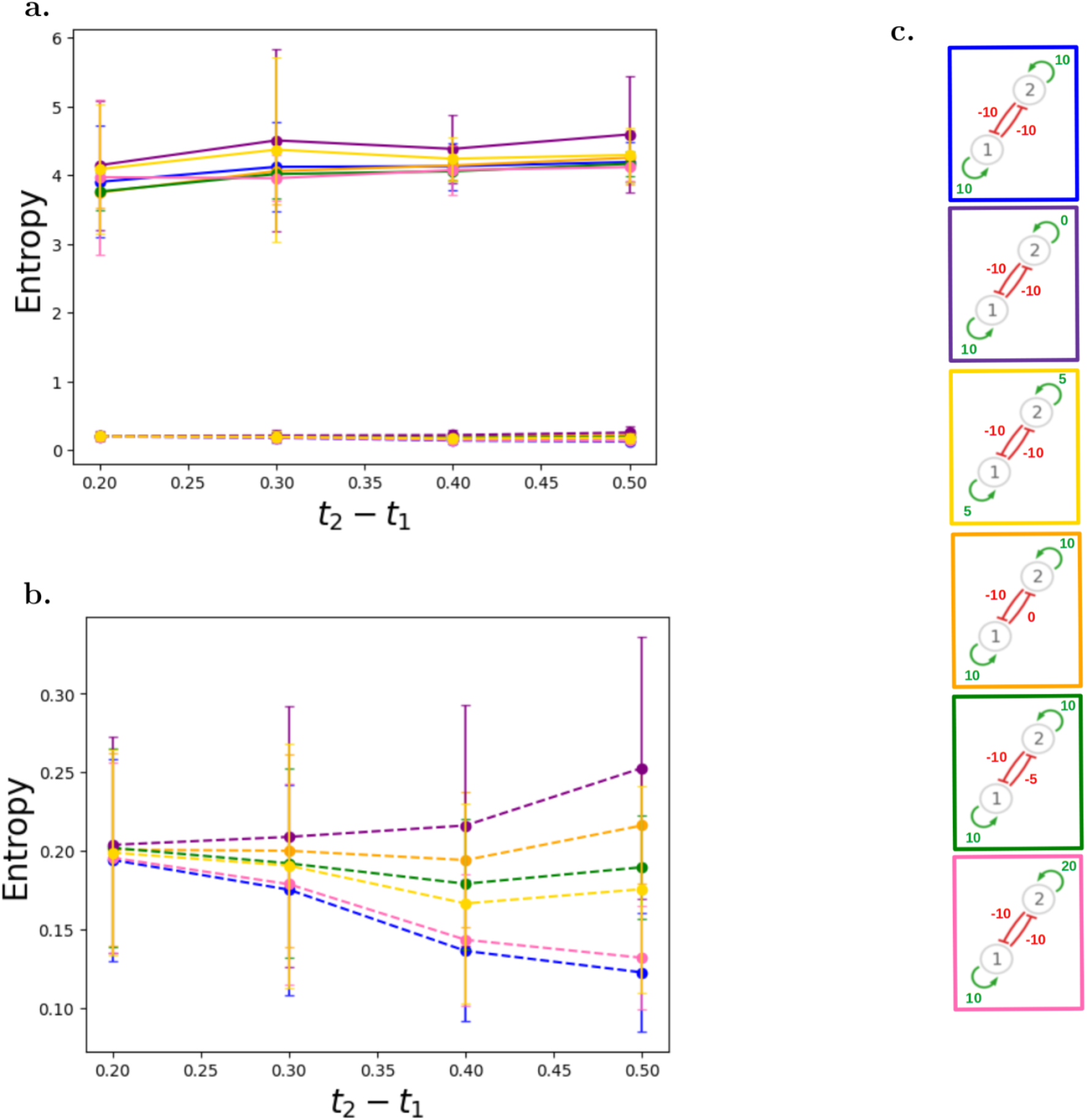
Entropy *H*_*D,B*_ as a function of the time difference *t*_2_ − *t*_1_ for different Bursty models, in transient state. **a**. Entropies *H*_*B*_ (dashed line) and *H*_*D,B*_ (straight line) means and standard deviations. Each color represents a set of interaction parameters of *B*_ref_ used to compute *H*_*D,B*_ (illustrated in Panel **c**). **b**. Zoom in on *H*_*B*_ values in Panel **a** for *t*_2_ − *t*_1_ ∈ [0.2,0.5]. **c**. Two-gene GRN used to compute *B*_ref_. Each scheme depicts interactions between gene 1 and gene 2, with green arrows representing activation (*θ* > 0) and red arrows representing inhibition (*θ* < 0). The blue-squared GRN is the toggle switch used in previous figures. Parameters values are detailed in Table 1.

**Figure 10:**
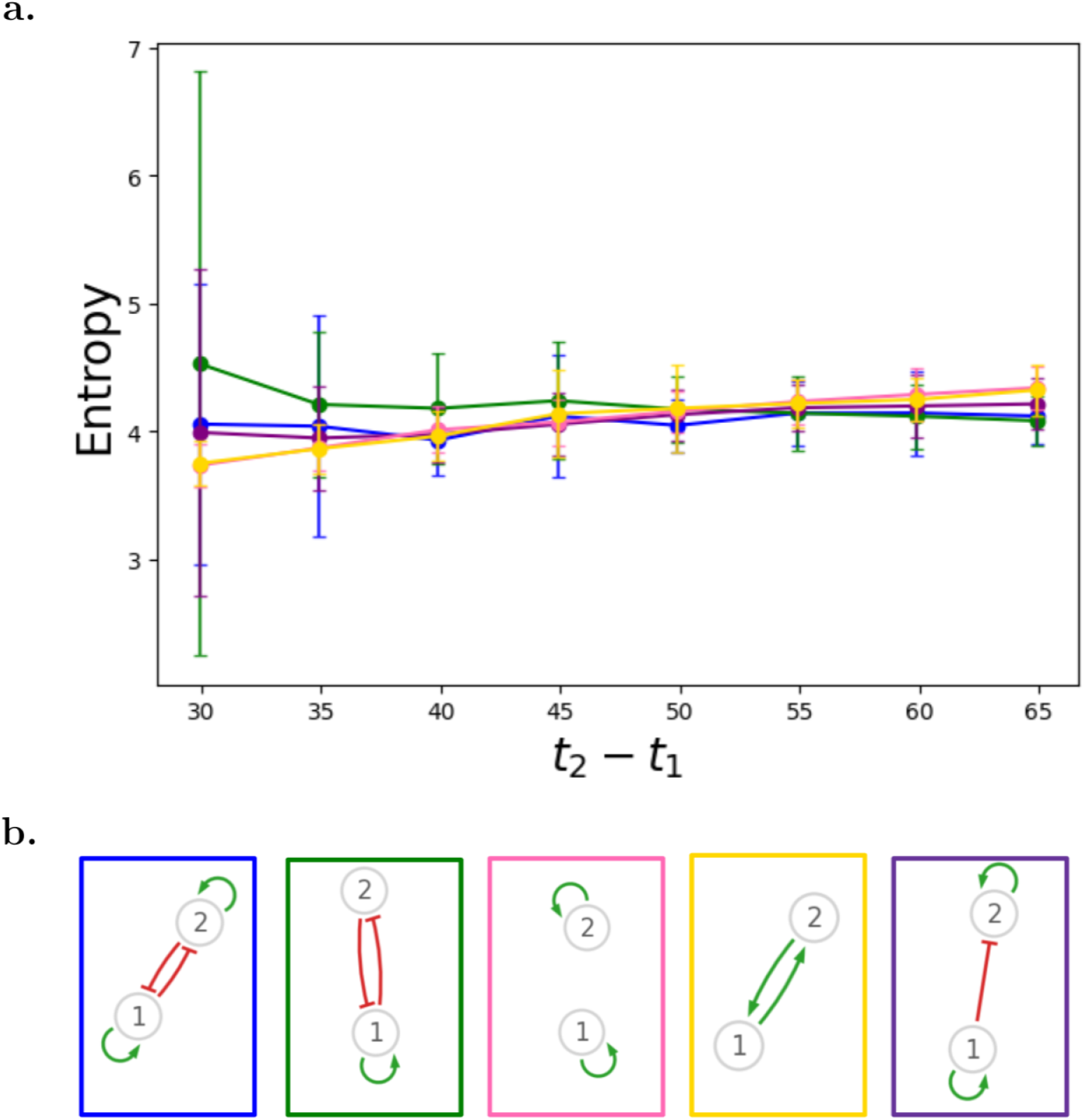
Entropy *H*_*D,B*_ as a function of the time difference *t*_2_ − *t*_1_ for various two-gene GRNs. **a**. Entropy *H*_*D,B*_ in transient state, mean and standard deviation. Each color represents a GRN introduced in Panel **b**.. Parameters are detailed in Table 1.**b**.Schematic representations of two-gene GRNs used in Panel**a**. Green arrows represent activations (*θ* > 0) and red arrows represent inhibitions (*θ* < 0).

**Figure 11:**
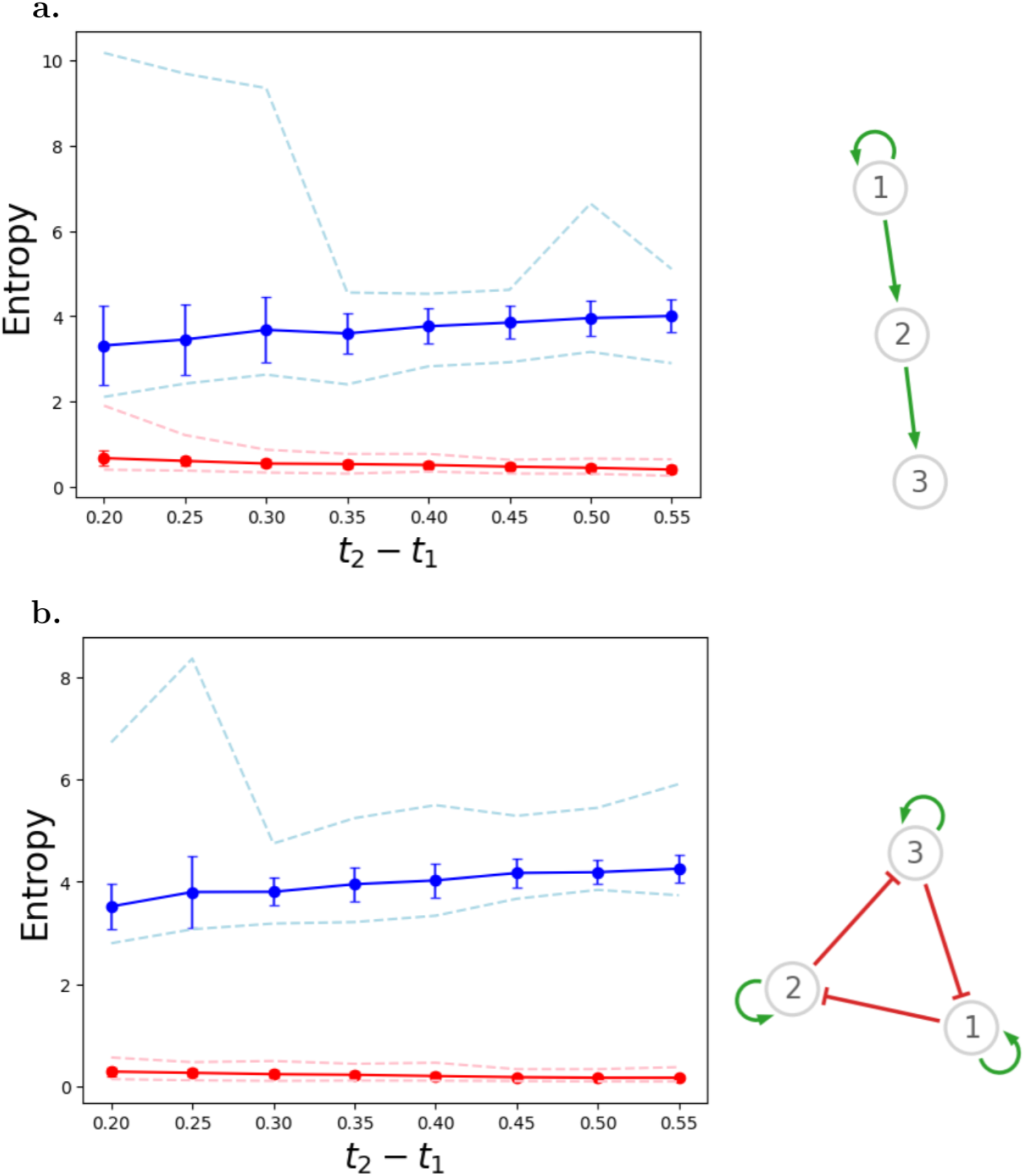
Entropy evolution as a function of the time difference *t*_2_ − *t*_1_ in the transient state for three-gene GRNs. **a**. *H*_*D,B*_ mean and standard deviations (blue line with error bars) as well as min and max values (dash blue lines) and *H*_*B*_ mean and standard deviation (red line with error bars) and min and max values (dash red lines), and the associated three-gene GRN. The schematic representation of the GRN highlights positive interactions (activation) between genes. **b**. *H*_*D,B*_ mean and standard deviations (blue line with error bars) as well as min and max values (dash blue lines) and *H*_*B*_ mean and standard deviation (red line with error bars) and min and max values (dash red lines), and the associated three-gene GRN (repressilator). The schematic representation of the GRN shows interactions between genes, with activations in green and inhibitions in red. Parameters are detailed in Table 1.

Overall, the Bursty model provides a framework to cell trajectory inference that performs better than standard diffusion.

### 3.3 Analysis for other GRNs

As illustrated in Figure 2, the toggle switch model of gene dynamics exhibits what can be considered as specific temporal dynamics. We subsequently challenged the Bursty process on other GRNs, first with various two-gene GRNs and then on three-gene GRNs. Results discussed hereafter and illustrated in Figures 10 and 11 have been obtained using various GRNs to compute both the distributions (*µ* and *ν*) and the reference coupling. This contrasts with the results shown in Figure 9, where only the reference coupling was modified by the GRNs, not the data.

Figure 10 shows the entropy *H*_*D,B*_ for different two-gene GRNs. For each GRN, the distributions and the Bursty process were computed from the corresponding Bursty model. The blue curve represents *H*_*D,B*_ values computed for the toggle switch. We observed that, whatever the two-gene GRN, dynamics of *H*_*D,B*_ were similar, quantitatively and qualitatively: it was mostly constant as the time difference *t*_2_ − *t*_1_ increased, and the variability was higher for small time differences.

Figure 11 shows the entropy *H*_*D,B*_ for two 3-gene GRNs. In the first one, gene 1 activates gene 2 which activates gene 3 (Figure 11a); in the second one, the so-called repressilator model, gene 1 inhibits gene 2, gene 2 inhibits gene 3, and gene 3 inhibits gene 1 (Figure 11b). Parameter values are the same than for the toggle switch (Table 1).

Figure 11a shows a similar range of variations for *H*_*D,B*_ and *H*_*B*_ as observed for the two-gene GRNs. Variations about the mean were, however, more important than for the two-gene GRNs, as highlighted by error bars and minimum and maximum values (dashed lines). The *H*_*D,B*_ curve remains well above the *H*_*B*_ curve, similarly to what has been obtained for two-gene GRNs. The same results held for the Repressilator model (Figure 11b), a standard three-gene GRN. Whatever the time difference, the mean of *H*_*D,B*_ was always higher than *H*_*B*_. For both GRNs, when *t*_2_ − *t*_1_ was small, the standard deviation of *H*_*D,B*_ was larger and the minimum and maximum values were widely separated. The two three-gene GRNs considered generated more differences between the Diffusive and the Bursty processes than the toggle switch model or any 2-gene GRN, as well as more variability.

In conclusion, for any two-gene or three-gene GRN, whatever the gene-gene interactions considered, the Bursty process always performed better than the standard Diffusive process in correctly transporting the distribution at time *t*_1_ into the distribution at time *t*_2._

## 4 Discussion

We explored the possibility that the Bursty model of gene dynamics would represent an alternative to the standard diffusion model used to infer trajectories in OT theory. Based on distributions of protein levels, the Bursty model proved indeed quite efficient in capturing cells trajectories more accurately than the Diffusive model (Figure 5). This would be expected if we considered cells moving around in a Waddington’s landscape [46] whose shape is constrained by the GRN structure [26, 39]. In this case, diffusion could not easily represent the variables acceleration and deceleration due to valleys and hills. Noticeably, we assumed for the sake of simplicity the same number of cells in both initial and final distributions, yet our entire analysis could be extended to the case with different cell numbers by normalizing the distributions.

When considering a toy model made of only one gene, we concluded that using the exact time-dependent distribution of the process was the best alternative to diffusion. Although expected, this result indicates that it is possible to very efficiently infer cell trajectories in a transcriptomic space. However, the exact distribution is only explicit for one gene. Extending the reference process *RB*_ref_ to GRN with multiple genes seems a very challenging task, probably impossible to do for complex gene interactions. An approximation of this distribution could be considered, yet the approximation might rapidly become coarse.

For a one-gene GRN, the computation of *RB*_ref_ and *D*_ref_ processes is almost instantaneous on a personal computer (64 GB RAM and 13th Gen Intel(R) Core(TM) i7-13700H), whereas the computation of *B*_ref_ takes approximately 7s (values averaged over 100 simulations). Increasing the number of genes does not impact the computational time for *D*_ref_, but it increases for *B*_ref_. The computational time is proportional to the number of genes in the GRN with almost no time-variability from one simulation to another, for two-gene up to six-gene GRNs where genes self-activate and inhibit the other genes (similar to the toggle switch model). For two genes, the computational time is 12s, for 3 genes, it is 13s, for 4 genes, 14s, for 5 genes 15.5s and for 6 genes 17s per simulation of the *B*_ref_ process, on a computational server (High-Throughput Computing plateform with 1 CPU and 2 GiB memories). This provides an optimistic perspective for increasing dimensionality.

As mentioned above, all investigations were performed on protein data, even though the Bursty model computes both mRNA and protein levels, which is necessary in order to represent transcriptional bursting and interactions between genes through their expression products (e.g., transcription factors). Although it would have been appropriate to focus our study on mRNA levels, as provided by scRNA-seq datasets, it is not possible to reduce the Bursty model to an mRNA-only model, due to the fact that mRNA are typically much less stable than proteins and thereby would not preserve the Markovian property of the GRN. This, however, makes it possible to reduce the Bursty model to a protein-only model that is well defined, along with quasi-steady-state distributions of mRNA levels conditionally on proteins [34, 37]. Moreover, Baradat and Ventre [47] showed that for such a model it is still possible to define and solve a Schrödinger problem. We therefore decided to focus on distributions of protein levels throughout this work. We nevertheless challenged the Bursty model against RNA data, similarly to what has been done by Schiebinger et al. [5]. Figure 12 shows that the *H*_*D,B*_ curve is higher than the *H*_*B*_ curve. Contrary to what was observed with protein level distributions, the *H*_*D,B*_ curve decreases as *t*_2_ − *t*_1_ increases, yet remaining far above the control curve. mRNAs are produced and degraded more rapidly than proteins, accelerating the approach to the stationary state. The same conclusions then hold, whether mRNA or protein data are used to infer cell trajectories.

**Figure 12:**
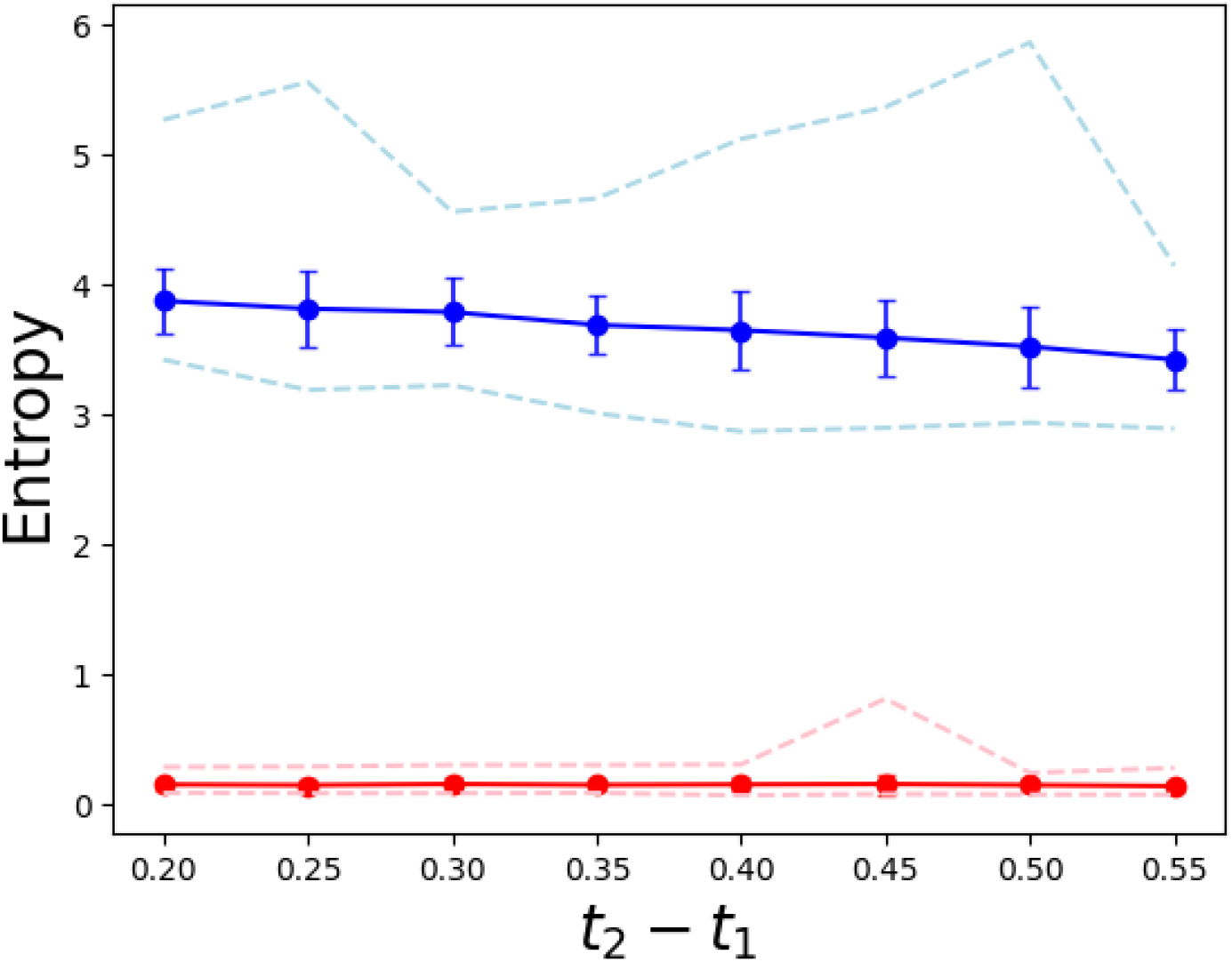
Entropies between reference processes for the toggle switch GRN for mRNA data. Mean and standard deviation (straight lines) of *H*_*D,B*_ (blue) and *H*_*B*_ (red) are displayed as functions of the time interval *t*_2_ − *t*_1_. Additionally, minimum and maximum values over 100 iterations are represented (dashed lines). Parameters are given in Table 1.

Therefore, the Bursty model represents a desirable alternative to diffusion for trajectory inference in OT, both when using protein data or mRNA data.

In order to use our approach on an experimental dataset, we would need first to infer the GRN that could be used to compute the reference process. For this, we could use CARDAMOM [36] which computes a Bursty-model GRN and the interaction parameter values from time-stamped scRNA-seq data. For now, degradation and synthesis rates have to be estimated before using the CARDAMOM algorithm but we are currently working on an extension allowing to fully calibrate the model from time-series of singlecell experimental data. Importantly, if the underlying GRN structure is not known, we know from Figure 9 that the method nevertheless correctly catches the trajectories for small GRNs. Although what impact it might have on larger GRNs remains to be explored, the results presented in this article thus pave the way to the extension of GRN inference algorithms to methods combining GRN and trajectory inference in an integrated framework.

## 5 Code Availability

All python codes are available at https://github.com/fourniecl/inference_trajectories_PDMP.

## 6 Acknowledgments

This work has received financial support from the CNRS through the MITI interdisciplinary programs through its exploratory research program (TROPIC project under Action 80PRIME 2023). This work was supported by the PEPR Santé Numérique under Project AI4scMED MultiScale AI for Single Cell-Based Precision Medicine (ANR-22-PESN-0002). We thank the computational center of IN2P3 (Villeurbanne/France), where part of the computations were performed.

## Supplementary File

## 1 Analytical solution for the reduced Bursty model

Lets consider the reduced Bursty model

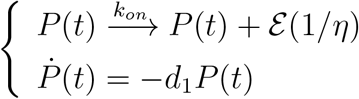

where *P*(*t*) is the protein level at time *t*, ℰ is the exponential distribution characterizing the random size of each burst (with mean value *η*), *k*_*on*_ is the burst rate, assumed to be constant, and *d*_1_ is the degradation rate of proteins.

We briefly explain how to derive the formula

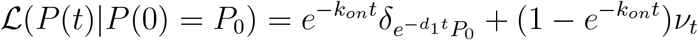

Where 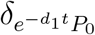 is the Dirac measure localized at 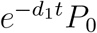, and 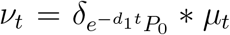 with *µ*_*t*_ the probability measure defined by the following density function:

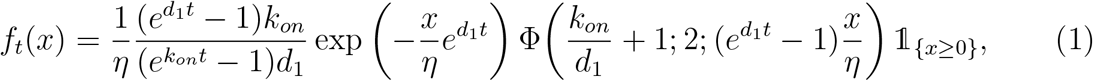

where Φ denotes the confluent hypergeometric (Kummer’s) function of the first kind.

Hence the probability measure *ν*_*t*_ has a density function *g*_*t*_(*x* | *P*_0_) given by

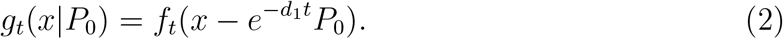

The corresponding Markov process (*X*_*t*_)_*t* ≥0_ (with values in ℝ _+_) is characterized by its infinitesimal generator *L* defined by [1]:

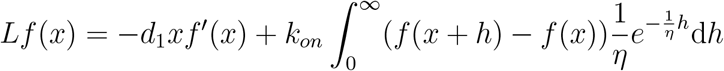

from which one can derive an evolution equation for the Laplace transform 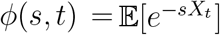, leading to the general solution [1]:

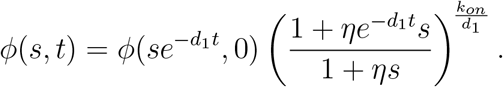

The first term in this product can be identified as the Laplace transform of 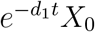, while the second term is the Laplace transform of *X*_*t*_ conditional on *X*_0_ = 0 (i.e., ℒ (*X*_*t*_ *X*_0_ = 0)). It turns out that the latter can be inverted explicitly using a classical series known as the confluent hypergeometric function of the first kind (denoted by *M*(*a*; *b*; *z*)), which is of practical interest since this special function is implemented in standard computing libraries such as SciPy. Manipulation of the series leads to the probability measure

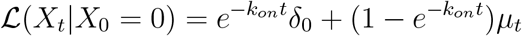

where *µ*_*t*_ has density function *f*_*t*_(*x*) given in (1), and the final expression (1)-(2) follows.

Interestingly, the weight of the singular part (Dirac measure at 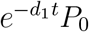 decreases exponentially with rate *k*_*on*_, in line with the property that in this model, a trajectory ceases to be deterministic precisely when the first burst occurs.

## 2 The stationary case

We studied the influence of the time difference *t*_2_ − *t*_1_ on the solutions of the Schrödinger problem when the system is in a stationary state. To do so, we defined *t*_1, *st*_ = 40 h and *t*_2, *st*_ ∈[42; 64] h. Results are displayed in Supplementary Figure 1.

Results are similar to the case of the transient state, except that the *H*_*D,B*_ curve decreases as *t*_2, *st*_ increases. This shows that despite the Diffusive model being less accurate than the Bursty model to infer trajectories, their performances become comparable when the final distribution becomes independent of the initial one, that is for large *t*_2, *st*_ values.

---

**Supplementary Figure 1:**
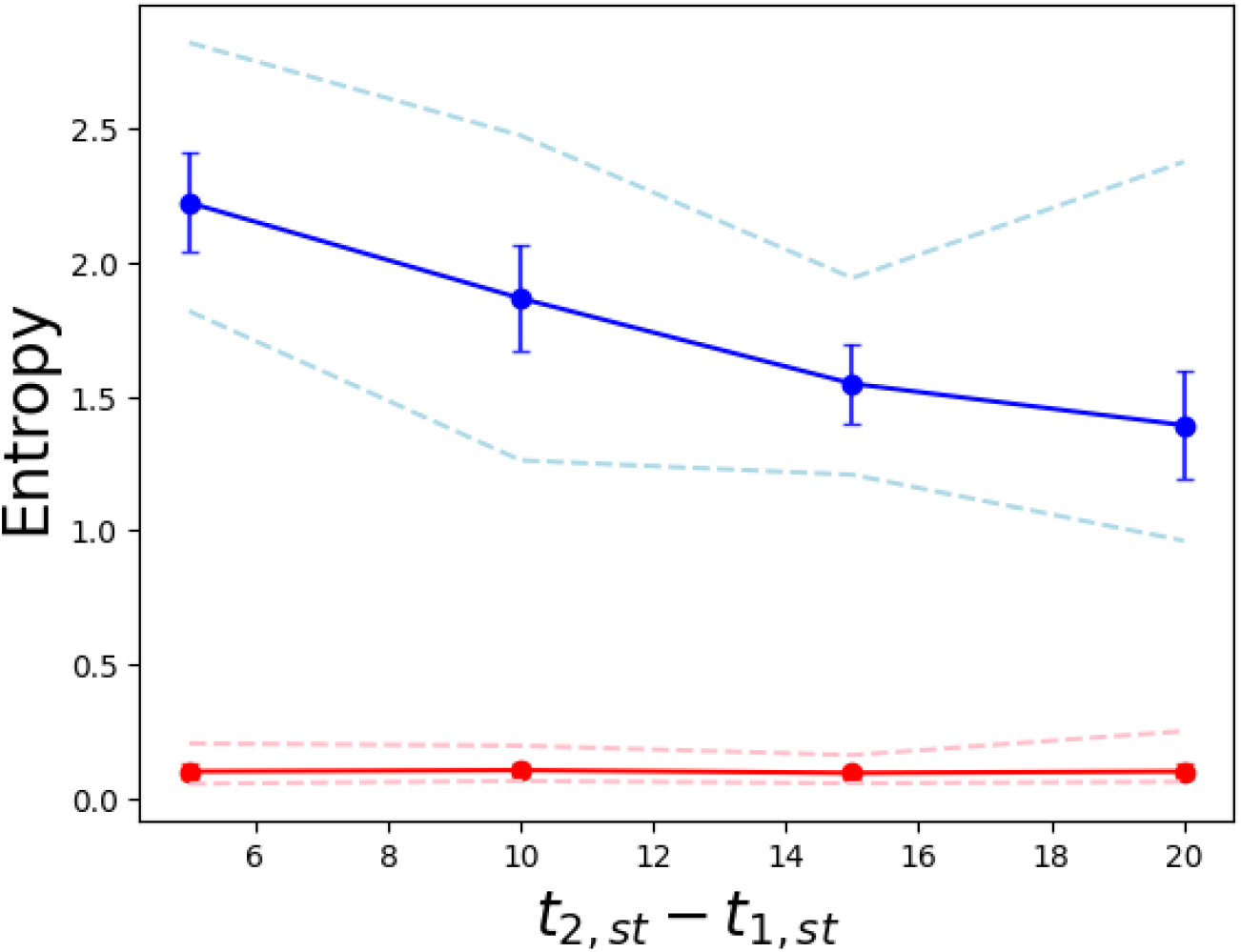
Entropies between reference processes for the toggle switch GRN, in the stationary state. Mean and standard deviation (straight lines) of *H*_*D,B*_ (blue) and *H*_*B*_ (red) are displayed as functions of the time interval *t*_2, *st*_ − *t*_1, *st*_. Additionally, minimum and maximum values over 100 iterations are represented (dashed lines). Other parameters are given in Table 1 (main text).

